# Bile Acid Regulates the Colonization and Dissemination of *Candida albicans* from the Gastrointestinal Tract by Controlling Host Defense System and Microbiota

**DOI:** 10.1101/2021.09.29.462497

**Authors:** Shankar Thangamani, Ross Monasky, Jung Keun Lee, Vijay Antharam, Harm HogenEsch, Tony Hazbun, Yan Jin, Haiwei Gu, Grace L. Guo

## Abstract

*Candida albicans* (**CA**), a commensal and opportunistic eukaryotic organism, frequently inhabits the gastrointestinal (GI) tract and causes life-threatening infections. Antibiotic-induced gut dysbiosis is a major risk factor for increased CA colonization and dissemination from the GI tract. We identified a significant increase of taurocholic acid **(TCA),** a major bile acid in antibiotic-treated mice susceptible to CA infection. *In vivo* findings indicate that administration of TCA through drinking water is sufficient to induce colonization and dissemination of CA in wild type and immunosuppressed mice. Treatment with TCA significantly reduced mRNA expression of immune genes *ang4* and *Cxcr3* in the colon. In addition, TCA significantly decreased the relative abundance of three culturable species of commensal bacteria, *Turicibacter sanguinis, Lactobacillus johnsonii*, and *Clostridium celatum*, in both cecal contents and mucosal scrapings from colon. Taken together, our results indicate that TCA promotes fungal colonization and dissemination of CA from the GI tract by controlling host defense system and intestinal microbiota that play a critical role in regulating CA in the intestine.

**Importance:** Broad-spectrum antibiotics, FDA-approved bile acid drugs, and probiotics used to control metabolic and infectious diseases profoundly alter the level of TCA in the gut. Furthermore, TCA level is highly altered in a subset of cancer, colitis and surgery patients who are highly susceptible to CA infection. Inadvertently, these therapies and disease conditions could be either promoting CA colonization and dissemination. Our findings indicate that TCA alone can induce fungal colonization and dissemination from the intestine. Results from this study will have a significant impact in understanding how bile acids interact with the microbiota and host in regulating invasive fungal infections that originate from the intestine and to develop potential new antifungal therapeutics.

## Introduction

*Candida albicans* (**CA**), a commensal and opportunistic eukaryotic organism, frequently inhabits the gastrointestinal (GI) tract and can cause life-threatening infections [1–3]. CA is present in small numbers in the healthy GI tract of humans; thus, CA is harmless in immunocompetent humans [4, 5]. While dysregulation in host defense system contribute to invasive CA infections, use of broad-spectrum antibiotics is a major predisposing risk factor for increased fungal colonization and subsequent dissemination from the intestine [6–13], CA is normally absent in the GI tract of adult mice; however, antibiotic treatment leads to CA colonization and dissemination in mice and closely resembles CA infection in human patients [4, 5, 13–21]. Because colonization of the GI tract is necessary for dissemination of CA and CA colonization resistance is nullified by antibiotic treatment, identifying the factors that play a critical role in CA colonization may pave the way to develop novel approaches to limit CA dissemination [5, 19].

Antibiotics alter the intestinal microbiota and lead to changes in the composition of the gut microbial metabolites that play a critical role in controlling several enteric bacterial pathogens [22–24], However, there is limited knowledge about the potential role of microbial metabolites in the regulation of CA colonization and pathogenesis. To address this gap in knowledge, we previously performed targeted metabolomics and identified several groups of metabolites that are differentially regulated in the gut contents of cefoperazone-treated CA-susceptible mice and control CA-resistant mice [25]. A bile acid, taurocholic acid (TCA), was identified as one of the major class of metabolites that was significantly increased in the antibiotic-treated mice [25]. Following synthesis in the liver, TCA enters the intestine and undergoes two major chemical modifications (*deconjugation and dihydroxylation*) carried out by gut microbes [26–28]. Antibiotic treatment depletes TCA-metabolizing commensal bacteria leading to increased levels of TCA in the gut contents [25]. Furthermore, the gut concentration of TCA is considerably altered in different diseases. Specifically, TCA is highly upregulated in immunocompromised cancer patients [29–31], drug-induced liver injury [32–34], high fat diets [35], and liver cirrhosis [36–41], conditions that are associated with severe morbidity and mortality caused by CA. Numerous FDA-approved drugs and probiotics currently used to treat various diseases also profoundly alter bile acid levels including TCA in the gut [42–51]. Therefore, understanding the role of TCA in the regulation of CA will gain insights into the bile-mediated regulation of gut fungi and will form a platform to develop novel therapies to control and treat invasive CA infections that arise as a result of antibiotic-induced dysbiosis and to control CA-associated infections in individuals with liver cirrhosis. Therefore, in this study, we investigated the role of TCA on CA colonization and dissemination from the intestine. Our findings indicate that TCA promotes fungal colonization and dissemination by altering intestinal defense system and microbiota.

## Materials and Methods

### Strains and Reagents

*Candida albicans* SC5314 was provided by Dr. Andrew Koh at the University of Texas Southwestern Medical Center [52]. This study used Yeast Peptone Dextrose (YPD) (242810, BD Difco) agar (BP1423, Fisher) as well as Bacteroides Bile Esculin Agar (BBE) (AS114, Anaerobe Systems) for various assays. Additionally, antibiotics kanamycin (J61272, Alfa Aesar), ampicillin (69-52-3, IBI Scientific), and streptomycin (S6501, Sigma-Aldrich) were used in media for fungal enumeration and cefoperazone (J65185, Alfa Aesar) was used in antibiotic pre-treatment. Taurocholic acid (TCA) (16215, Cayman Chemicals) and cyclophosphamide (PHR1404, Millipore Sigma) were purchased for these studies. Additional kits and reagents used included: hydrogen sulfide gas assay (ZAN-5084, Cell Biolabs), fluorescein isothiocyanate-dextran (FITC-dextran) (FD4, Millipore Sigma), taurine (A12403, Alfa Aesar), and sodium formate (A17813, Alfa Aesar).

### Mouse studies

Male and female C57BL/6J mice (000664) and BALB/cJ mice (000651) of ages 6-12 weeks were purchased from the Jackson Laboratory. All animal studies conducted were approved by the Institutional Animal Care and Use Committee (IACUC) at Midwestern University under MWU IACUC Protocol #4014.

### Fungal colonization and dissemination in the immunosuppressed mouse model

Studies specifically examining survival and dissemination of mice infected with *C. albicans* SC5314 under immunosuppressive conditions were performed with BALB/c mice. Male and female mice (age 8 to 12 weeks old) were treated with or without cefoperazone (0.5mg/mL) via drinking water, and drinking water was replaced with fresh antibiotic water every two days. After 7 days, all mice were infected with *Candida albicans* SC5314 via oral gavage with 1-2 x 10^8^ CFU/mouse. Three days post infection, all mice were treated with 150 mg of cyclophosphamide (cyclo) per kilogram of mouse body weight by intraperitoneal injection. A second and third dose of cyclophosphamide were given again at 5- and 7-days post infection, respectively. Mice were monitored for survival, and euthanized if mice were moribund. After euthanasia, cecal contents, liver, and kidneys were collected and processed for fungal burden to assess the fungal dissemination as described before [25]. Briefly, gut contents and tissue homogenates were serially diluted and plated onto YPD agar plates containing broad spectrum antibiotics and incubated at 30°C for 24 hours to determine the colony forming unites (CFU).

### Effect of TCA on fungal colonization and dissemination in mice

Male and female wild type C57B/6 mice were infected with *C. albicans* SC5314 via oral gavage with 1-2 x 10^8^ CFU/mouse. One group of mice was treated with 1% TCA in the drinking water and the water was changed every day. Control groups received sterile water. Fecal contents were collected at the indicated time points to determine the fungal load in feces. Mice were monitered for survival and cecal contents, liver, and kidneys were processed to determine the fungal load and dissemination as described before [25]. To determine the effect of TCA on fungal colonization and dissemination in the immunosuppressed mouse model, male and female BALB/cJ mice (age 10 to 12 weeks old) were infected as described above and all mice were treated with 150 mg of cyclophosphamide (cyclo) per kilogram of mouse body weight through intraperitoneal injection. Second and third dose of cyclophosphamide were given again at 5- and 7-days post-infection respectively.

### FITC-Dextran permeability assay

Gut permeability was measured in infected mice using a FITC-dextran assay. Wild type C57BL/6 mice were infected with *Candida albicans* SC5314 via oral gavage with 1-2 x 10^8^ CFU/mouse and treated with or without 1% TCA through drinking water. After 10 days post-infection, mice were administered orally 150 uL PBS containing 15mg FITC-dextran. The mice were anesthetized four hours later and blood was collected via retro-orbital route. Blood samples were then centrifuged to collect serum. Serum samples were processed via a 2-fold serial dilution in a 96-well plate and fluorescence was measure via plate reader (excitation: 485nm; emission: 520nm). Standard curves were made using serially diluted FITC-Dextran in PBS to determine the FITC-Dextran levels in serum samples.

### RNA sequencing and analysis

Wild type C57BL/6 mice were infected with *C. albicans* SC5314 via oral gavage with 1-2 x 10^8^ CFU/mouse and treated with or without 1% TCA through drinking water. At 10 days post-infection, colon was collected and flash frozen in liquid nitrogen. Total RNA was collected using Zymo Research RNA kit as per the manufacturer’s instructions. Total RNA-Seq libraries were constructed from 500 ng of total RNA and rRNA was removed. Libraries were prepared using the Zymo-Seq RiboFree Total RNA Library Prep Kit (Cat # R3000) according to the manufacturer’s instructions. RNA-Seq libraries were sequenced on an Illumina NovaSeq to a sequencing depth of at least 30 million read pairs (150 bp paired-end sequencing) per sample. Sequence Data Alignments and Differential Expression Analysis: NovaSeq paired-end 150-bp reads from Total RNA-Seq data files were first adaptor trimmed, and then analyzed using the STAR program (version 2.6.1d) for alignment of short reads to the genome of interest. Transcript counts were inferred from alignment files. All transcripts with either 0 or 1 counts were removed. Gene expression was measured using EdgeR. The DESeq2 R package was used for differential expression analysis using the gene feature.

### Microbiome sequencing and analysis

Wild type C57BL/6 mice were infected with *C. albicans* SC5314 via oral gavage with 1-2 x 10^8^ CFU/mouse and treated with or without 1% TCA through drinking water. After 10 days post-infection, cecal contents and mucosal scrapings from colon was collected using glass slides to determine the microbiome composition. Microbiome sequencing was carried out at Zymo Research corporation (Irvine, CA). The ZymoBIOMICS® Microbial Community Standard (Zymo Research, Irvine, CA) was used as a positive control for each DNA extraction, if performed. The final library was sequenced on Illumina® MiSeqTM with a v3 reagent kit (600 cycles) and the relative abundance of bacteria were analyzed as described elsewhere [25].

### Immunofluorescence staining

Colon from untreated and TCA treated mice infected with CA were collected after 10-12 days of infection. Colon tissue was fixed in 10% neutral buffered formalin solution and sectioned at 5 μm, and stained with DAPI and ZO-1 Antibody (21773-1-AP, Thermofischer). Stained tissues were imaged (40X) using a Keyence BZ-X700 microscope.

### Metabolomics

Bile acid metabolites (TCA and DCA) levels in the cecal contents were determined as described previously [25].

### Statistical analysis

Statistical analyses were performed using GraphPad Prism 6.0 (Graph Pad Software, La Jolla, CA). *P* values were calculated using the Mann-Whitney U test or unpaired Student’s t-test or one way-ANOVA followed by Bonferroni comparison or Kaplan-Meier (log rank) survival test as indicated. *P* values of (*≤ 0.05) (**P≤ 0.01) (***p≤ 0.001) (****p≤ 0.0001) were considered as significant.

## Results

### TCA is the major bile acid metabolite up-regulated in the cefoperazone-treated mice susceptible to CA infection

To identify the metabolites that regulate CA colonization and dissemination, we treated BALB/c mice with or without a broad-spectrum antibiotic (cefoperazone), orally infected them with CA, and then injected them with cyclophosphamide intraperitoneally. As expected, all mice in the antibiotic-treated group died within 6 days of infection (**Fig. 1A**). Furthermore, the fungal load was significantly higher in the cecal contents and liver of antibiotic-treated mice, thus confirming increased CA colonization and dissemination from the GI tract of antibiotic-treated mice compared to control groups (**Fig. 1B**). Previous findings from our lab indicates that more than 200 metabolites were differentially regulated in the cecal contents of control and antibiotic-treated mice [25]. Bile acids are one of the major metabolites differentially altered in the CA-susceptible mice compared to control groups [25]. Among bile acid metabolites, our results indicate that six bile acids were significantly upregulated in the antibiotic-treated CA susceptible mice (**Fig. 1C-D**). Interestingly, only the abundance of TCA was significantly higher than all other bile acid metabolites that were increased in the antibiotic-treated CA susceptible mice compared to untreated control groups (**Fig. 1D**). These results indicate that TCA is the highly upregulated bile acid in the antibiotic-treated CA-susceptible mice compared to control groups.

**Figure 1.**
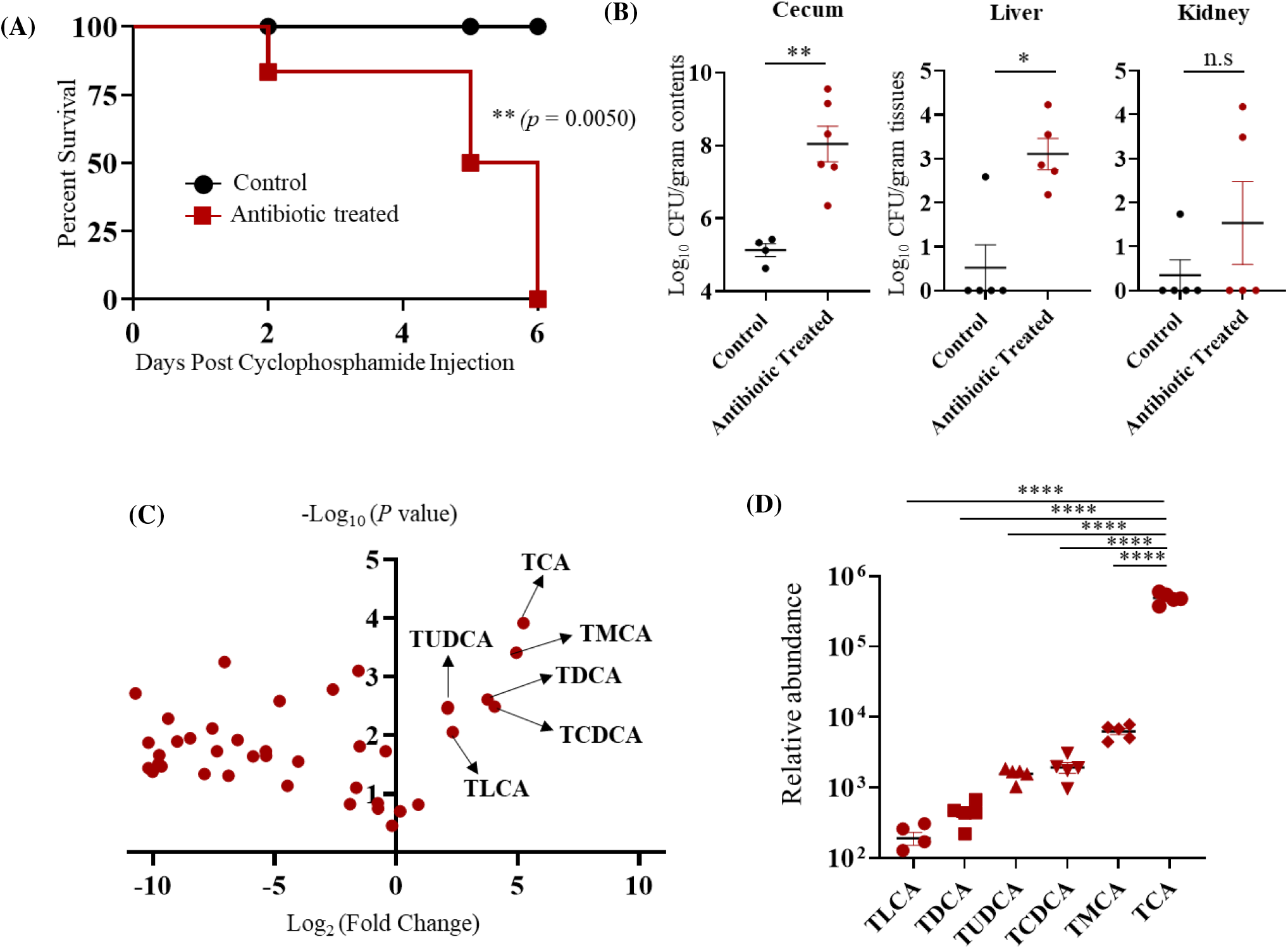
Taurocholic acid (TCA) is the major bile acid metabolite up-regulated in the cefoperazone-treated mice susceptible to CA infection. **(A)** BALB/c mice were fed sterile water in the presence or absence of cefoperazone (0.5 mg/mL) for 7 days and then infected with ~2 × 10^8^ CFU CA SC5314 via oral gavage. Antibiotic treatment was continued until the end of the experiment. Three days post-infection, all mice were injected with three doses of cyclophosphamide intraperitoneally (150mg/kg body weight) and monitored for survival. A log rank test was performed using 95% confidence intervals; statistical significance was calculated to compare the antibiotic-treated and untreated control groups. **(B)** Fungal load from cecum, liver and kidney were collected immediately after death in the antibiotic-treated groups and mice euthanized after 6 days post-cyclophosphamide treatment from control groups. Statistical significance was evaluated using the Mann-Whitney U test. **(C)** Relative Log2 fold-change of bile acid metabolites in antibiotic-treated C57BL/6 mice relative to control groups. **(D)** Relative abundance of bile acid metabolites that were highly upregulated and significant in the antibiotic-treated group relative to control groups. At least five mice per group was used and the data represents mean ± SEM. one way-ANOVA followed by a multiple comparison using Bonferroni correction. *P* values of (*≤ 0.05) (**P≤ 0.01) (***p≤ 0.001) (****p≤ 0.0001) were considered as significant.

### TCA alone induces fungal colonization and dissemination from the GI tract in the absence of antibiotics and immunosuppressive agents

To dissect the specific role of TCA in regulating CA colonization and dissemination *in vivo*, C57BL/6J mice infected with CA were treated orally with or without 1% TCA in the drinking water (**Fig. 2A)**. The fungal load in stool samples and mortality were monitored. Mice treated with TCA alone died 15 days post-infection (**Fig. 2B)**. Fungal load in the feces of the TCA group was significantly higher starting day 4 post-infection compared to the untreated control groups (**Fig. 2C)**. Furthermore, the TCA-treated group had significantly higher fungal loads in the cecal content, liver, and kidney, whereas no CA was detected in the liver and kidney from the untreated mice (**Fig. 2D)**. These results indicate that TCA alone can induce fungal colonization and dissemination from the GI tract in the absence of antibiotics and (or) immunosuppression.

**Figure 2.**
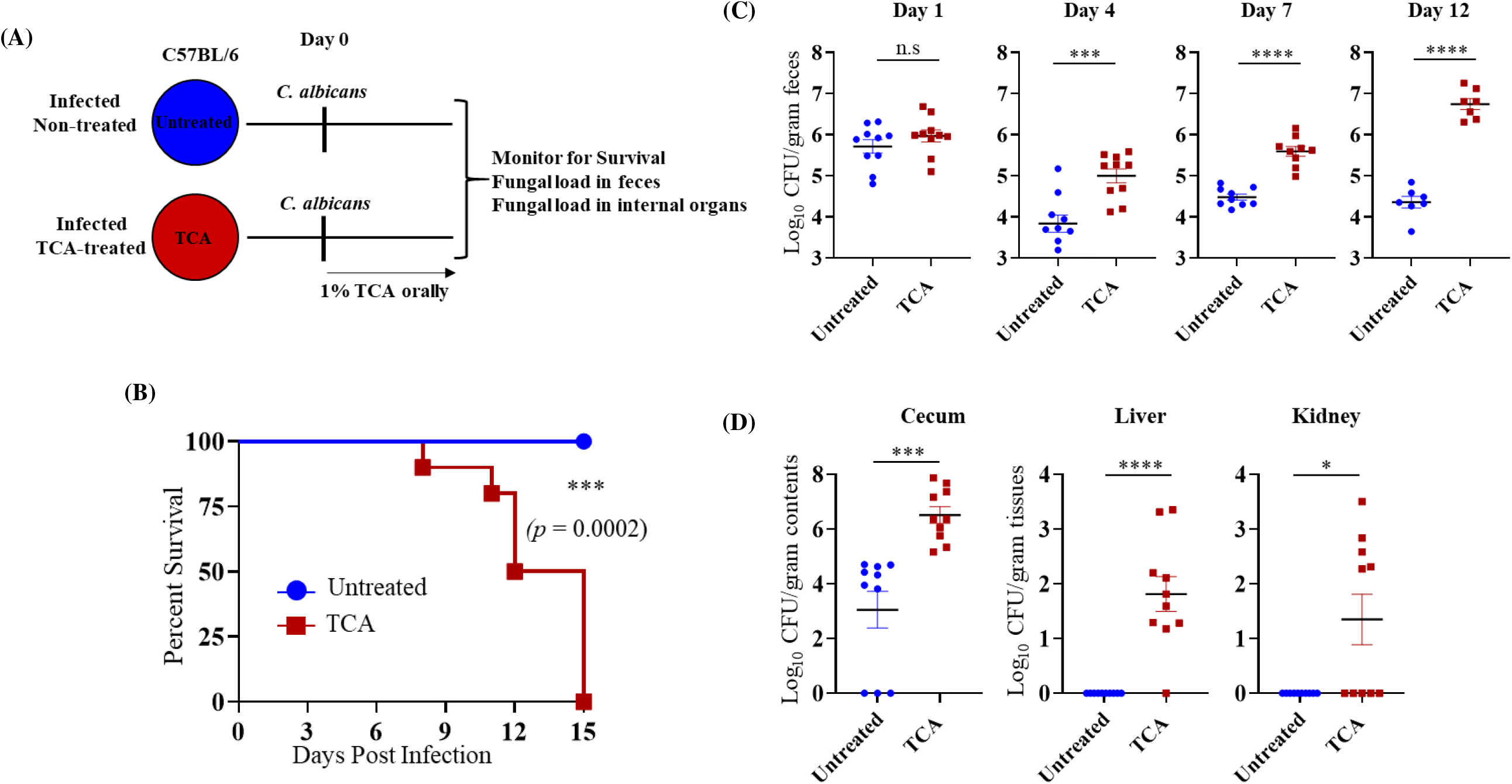
TCA alone induces fungal colonization and dissemination in the absence of antibiotics and immunosuppression. **(A)** Experimental outline. C57BL/6 mice infected with ~2 × 10^8^ CFU CA SC5314 via oral gavage. Untreated mice received sterile drinking water (untreated group); treatment group received sterile water containing 1% TCA (TCA group). **(B)** Mice were monitored for survival. A log rank test was performed using 95% confidence intervals; statistical significance was calculated to compare the antibiotic-treated and untreated control groups. **(C)** Fungal load from feces collected from untreated and TCA groups 1-, 4-, 7-, and 12-days post-infection. **(D)** Fungal load from cecum, liver, and kidney from dead mice collected immediately in the TCA-treated groups and mice euthanized 15 days post-infection for untreated groups. Ten mice per group was used and the data represents mean ± SEM. Statistical significance was evaluated using the Mann-Whitney U test. *p* values ≤0.05 (*) or ≤0.0l (**) were considered statistically significant.

### TCA induces fungal colonization and dissemination from the GI tract of immunosuppressed mice in the absence of antibiotic treatment

Since antibiotic induced gut dysbiosis is necessary to induce CA colonization and dissemination in immunosuppressed mice (**Fig. 1A**) [5], we tested whether administration of 1% TCA in the drinking water in immunosuppressed mice can promote fungal colonization and dissemination **(Fig. 3A)**. Interestingly, BALB/c mice that received TCA started to die 6 days post-cyclophosphamide injection and all the mice were dead by day 9 after the first dose of cyclophosphamide. On the other hand, the mice that were infected with CA and received three doses of cyclophosphamide survived **(Fig. 3B).** The fungal load was significantly increased in cecal contents and liver of the TCA treated group compared to immunosuppressed mice that received only sterile drinking water **(Fig. 3C)**. These results suggest that an increased concentration of TCA in the intestine promotes fungal colonization, dissemination and mortality even in the absence of antibiotic treatment in immunosuppressed mice.

**Figure 3.**
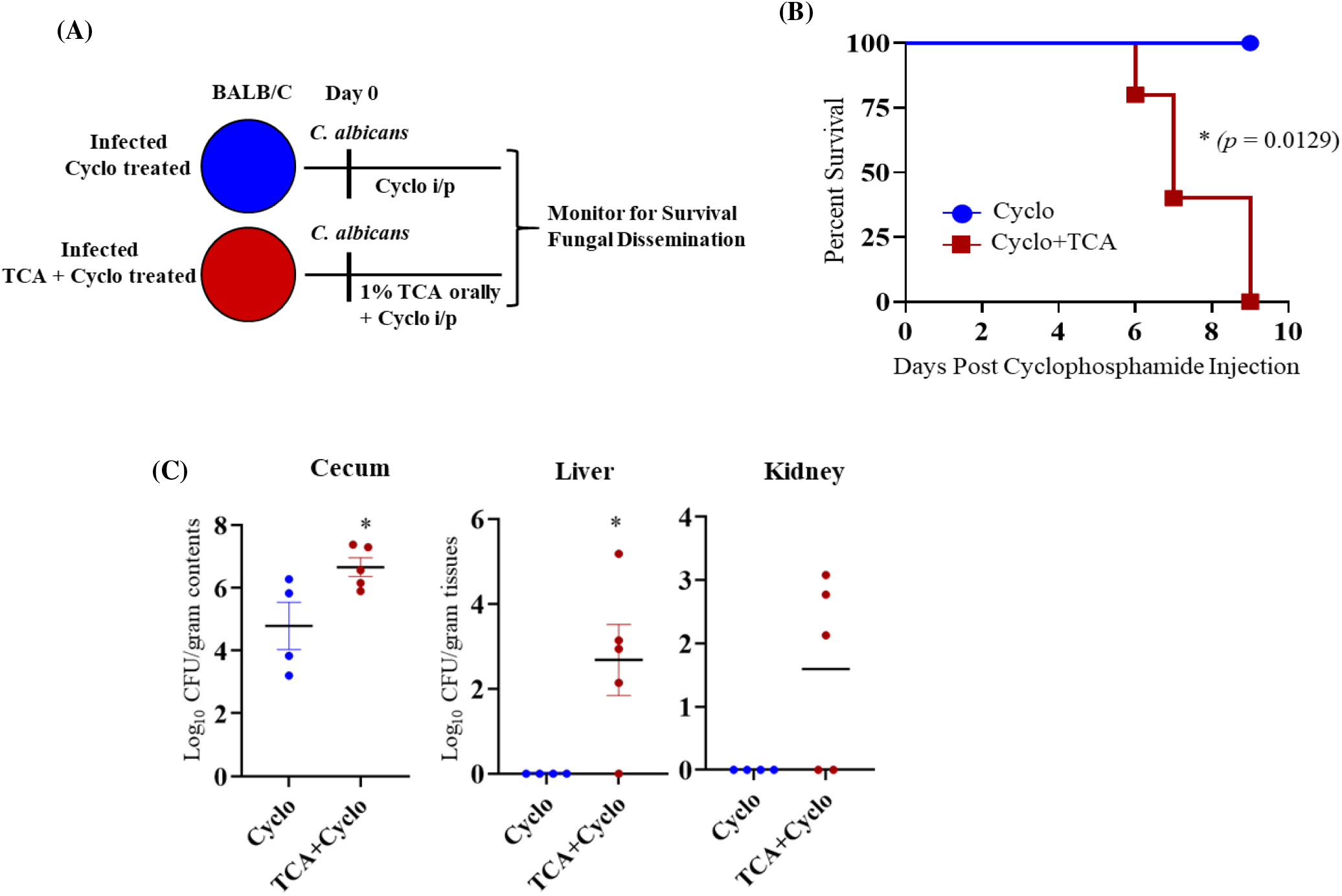
TCA induces fungal dissemination from the GI tract of immunosuppressed mice in the absence of antibiotic treatment. **(A)** Experimental outline. BALB/c mice infected with ~2 × 10^8^ CFU CA SC5314 via oral gavage received sterile water with or without containing 1% TCA. Three days post-infection, all mice were injected with three doses of cyclophosphamide intraperitoneally (150mg/kg body weight). **(B)** Mice were monitored for survival and log rank test was performed using 95% confidence intervals; statistical significance was calculated to compare the antibiotic-treated and untreated control groups. **(C)** Fungal load from cecum, liver, and kidney collected immediately after death in the TCA+ cyclo treated groups and mice euthanized 9 days post-infection for cyclo groups. 4-5 mice per group was used and the data represents mean ± SEM. Statistical significance was evaluated using the Mann-Whitney U test. *p* values ≤0.05 (*) or ≤0.0l (**) were considered statistically significant.

### TCA enhanced intestinal permeability and reduced expression of a tight junction protein

The intestinal barrier function is critical to prevent enteric pathogens from disseminating from the GI tract to systemic organs [5, 53]. Since TCA treatment induced CA dissemination from the GI tract, we evaluated if TCA dysregulated the intestinal barrier function with a FITC-Dextran permeability assay. The TCA-treated group had significantly increased levels of FITC-Dextran in the blood compared to the untreated control groups (**Fig. 4A**). Furthermore, expression of the ZO-1 tight junction protein was decreased in the colon of TCA-treated mice (**Fig. 4B**). Altogether, these results indicate that TCA dysregulates the intestinal barrier function which may contribute to CA dissemination from the intestinal tract.

**Figure 4.**
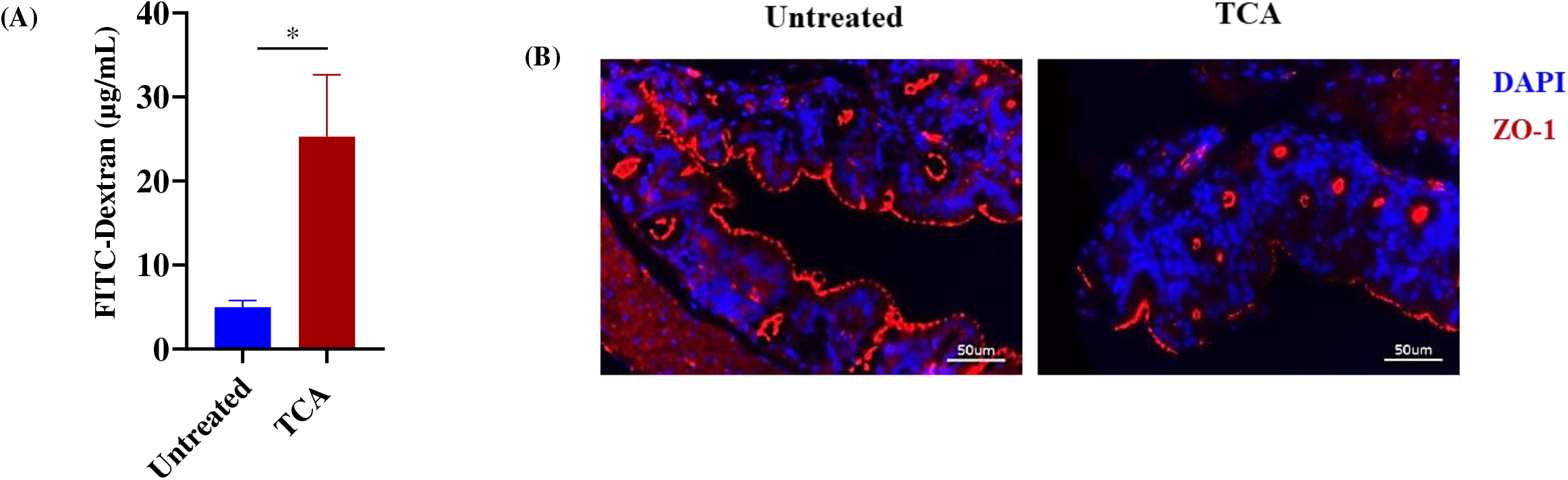
TCA increases the intestinal permeability leading to fungal dissemination from the GI tract. **(A)** Gut permeability was measured in infected mice using a FITC-dextran assay. At 10 days post-infection, male and female BL57/6J mice in both control and TCA-treated groups infected with CA were given an oral gavage of 150 μL PBS containing 15 mg FITC-dextran. Four hours after administering FITC-dextran, mice were anesthetized, and blood was collected via retro-orbital bleed. Blood samples were processed via a two-fold serial dilution in a 96-well plate and fluorescence was measure via a plate reader (excitation: 485 nm; emission: 520 nm). Data represent mean ± SEM. Data represent mean ± SEM. Statistical significance was evaluated using the Mann-Whitney U test. *p* values ≤0.05 (*) or ≤0.0l (**) were considered statistically significant. **(B)** ZO-1 tight junction protein expression in untreated and TCA-treated mice. Colon tissue from untreated and TCA-treated mice from CA-infected mice (10 days post-infection) were stained with ZO-1 and DAPI antibodies. Representative images are shown here.

### TCA downregulates *ang4* and *Cx3cr1* expression in the colon tissue

CA can cause severe invasive disease and mortality once critical components of the host defense system are compromised [5, 53–56]. To identify if TCA dysregulates the intestinal host defense system to induce CA colonization and dissemination, colonic tissue from untreated and TCA-treated mice collected after 10 days of CA infection and treatment were RNA sequenced to examine the expression of host defense genes (**Fig. 5A)**. Among the host defense genes examined, the relative expression of angiogenin 4 (*ang4*), an antimicrobial peptide, and CX3CR1 (*Cx3cr1*), a chemokine receptor, was significantly downregulated in TCA-treated mice compared to untreated mice infected with CA (**Fig. 5A and 5B**). Bile acids receptors play an important role in bile acid signaling [49, 57–61]. Further, our findings indicate that among the receptors examined, the relative expression of *Slc10a2* and *Nr1h4* that encode apical sodium-dependent bile acid transporter (ASBT) and farnesoid X receptor (FXR) receptors, respectively, was increased in the TCA-treated mice compared to untreated groups infected with CA (**Fig. 5C)**.

**Figure 5.**
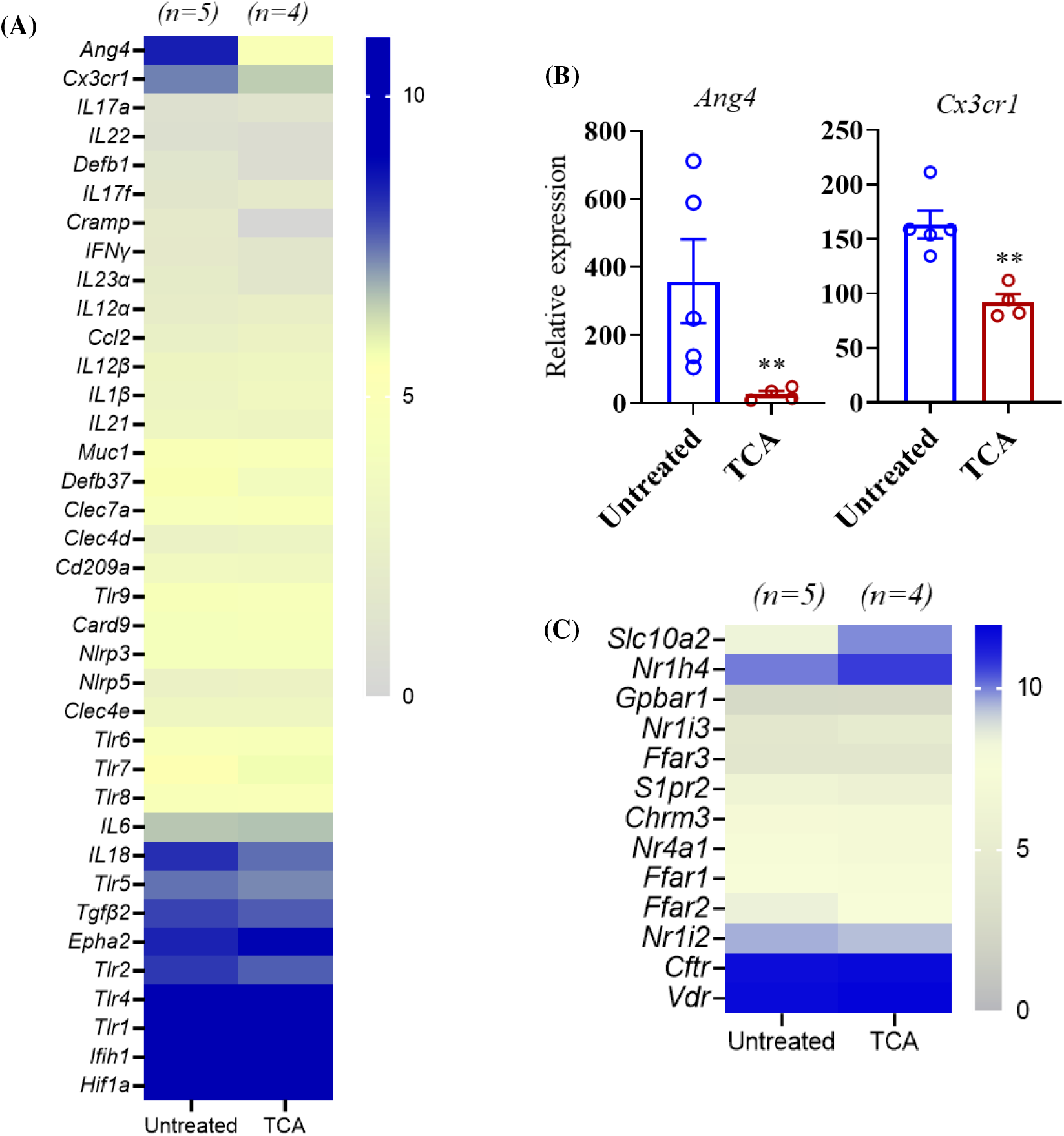
TCA inhibits the expression of *ang4* and CX3CR1 in the colon *in vivo.* Colon tissue from untreated and TCA-treated mice collected after 10 days of CA infection and treatment were RNA sequenced to examine the expression level of host defense genes. (**A**) Average relative expression of host defense genes in untreated and TCA treated mice were shown. (**B**) Relative expression of *ang4* and *Cx3cr1* in untreated and TCA treated mice were shown. (**C**) Average relative expression of intestinal metabolite receptors in untreated and TCA treated mice were shown. Four to five mice per group was used. Data represent mean ± SEM. Statistical significance was evaluated using the Student’s t test.

### TCA alters microbial composition in both luminal and mucosal parts of the GI tract

Next, we examined if TCA changes the composition of intestinal microbiota, as commensal bacteria are also essential to provide colonization resistance to CA [5, 25]. We analyzed the intestinal microbiota composition from the cecal contents and mucosal scrapings from colon in the untreated control and TCA treated mice infected with CA. Our results revealed that several culturable and unculturable bacterial members were significantly altered in cecal contents and colon mucosal scrapings in both groups (**Supplementary Figures 1-4**, **Fig. 6A and 6B**). Out of the top 20 statistically significant culturable and unculturable bacterial members found in cecal contents and colon mucosal scrapings, we focused on culturable bacterial members whose 16s reads could be mapped at species-level resolution. The relative abundance of three culturable bacterial members: *Turicibacter sanguinis* (cecum: 5.2% versus 0.90%; colonic mucosa: 5.2% versus 0.65%), *Lactobacillus johnsonii* (cecum: 26.4% versus 8.0%; colonic mucosa: 24.6% versus 6.8%), and *Clostridium celatum* (cecum: 5.3% versus 0.76%; colonic mucosa: 4.3% versus 0.44%) were significantly decreased in both cecal content and colon scrapings from TCA-treated mice compared to untreated mice (**Fig. 6C and 6D**). In mucosal scrapings, *L. johnsonii* constituted close to 25% of 16s reads but was three times more depleted in mice fed with TCA-infused drinking water. Comparatively, the diminishment as a result of TCA was 6-7-fold for reads mapping to *T. sanguinis* and *C. celatum.* Taken together, the relative abundance of three culturable bacterial members (*T. sanguinis, L. johnsonii*, and *C. celatum*) were significantly decreased in both cecal content and colon scrapings from TCA-treated mice compared to untreated mice infected with CA.

**Figure 6.**
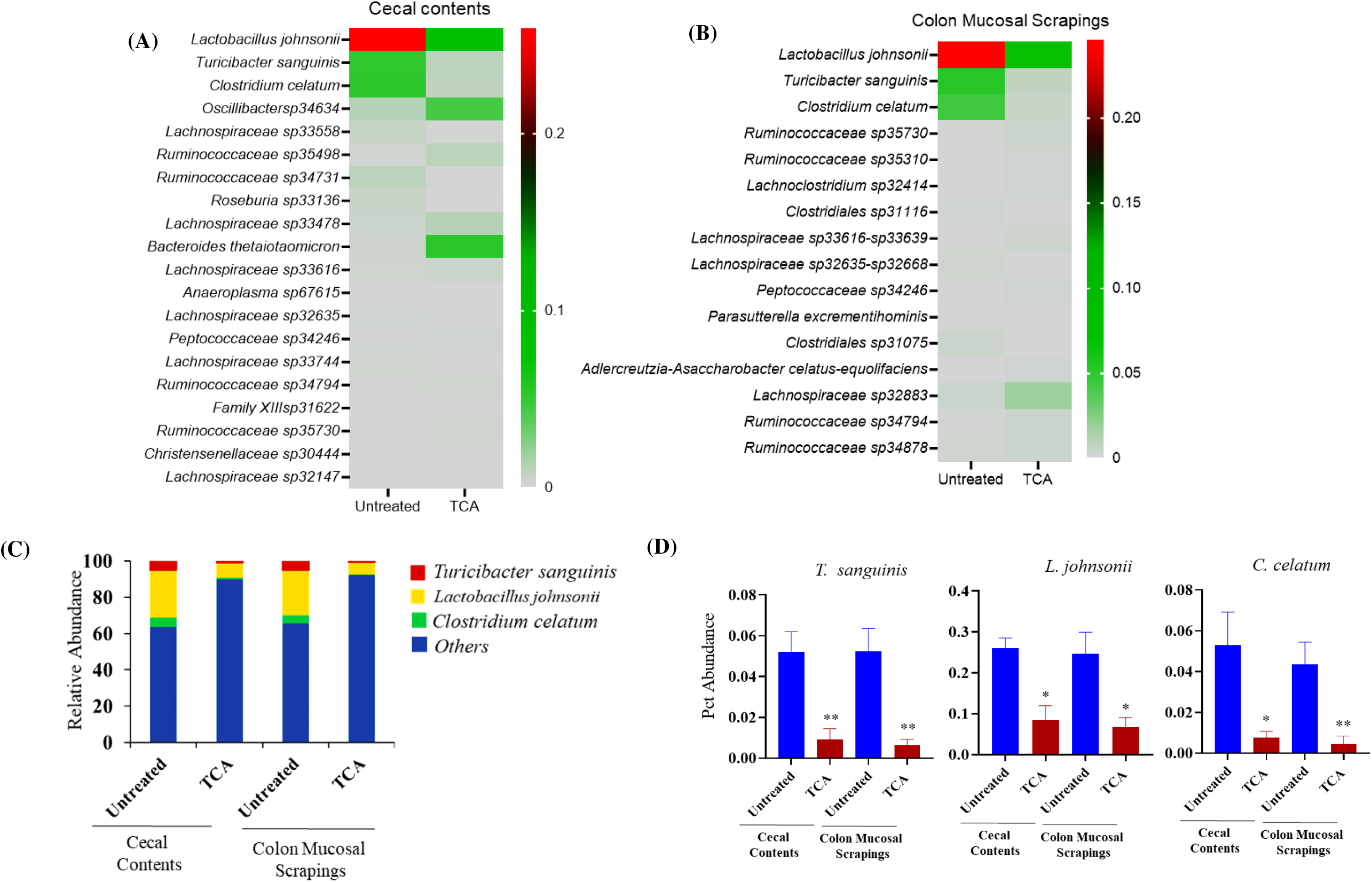
TCA alters the relative abundance of intestinal microbiota. (**A** and **B)** C57BL/6 mice were infected with ~2 × 10^8^ CFU CA SC5314 via oral gavage. Control mice received sterile drinking water (untreated group); the treatment group received sterile water containing 1% TCA (TCA group). Ten days post-infection, cecal contents and colon scrapings were collected to determine the relative abundance of intestinal microbiota. Bacterial members that are significantly altered in the TCA group compared to the untreated control groups both in cecal content and colon mucosal scrapings were shown here. Five mice per group was used. Data represent mean ± SEM. Statistical significance was evaluated using the Student’s t test.

## Discussion

Our findings indicate that TCA is one of the most strongly upregulated major bile acids in the antibiotic-treated mice susceptible to CA infection. After synthesis in the liver, TCA enters the small intestine and undergoes deconjugation and dihydroxylation, which are carried out by bile salt hydrolases and bile acid inducible enzymes present in gut microbes. These two chemical modifications convert TCA to DCA in the cecum [47, 57, 62–67]. Treatment with antibiotics decreases the microbial metabolism of TCA and induces an increase of this bile acid. Treatment of wild type and immunosuppressed mice with TCA alone promoted fungal colonization and dissemination from the GI tract in the absence of antibiotic treatment. Our studies revealed three primary mechanisms that underlie the TCA-induced enhanced CA colonization and dissemination from the GI tract including: (i) deficiencies in host defense mechanisms, (ii) disruption of the commensal microbiota which permits intestinal overgrowth of CA and (iii) intestinal barrier dysfunction [5, 53].

TCA significantly decreased the expression of *ang4* and *Cx3cr1* in the colon. Interestingly, both angiogenin-4 and CX3CR1 are involved in antifungal defense [55, 68]. Mice have three orthologous *ang* antimicrobial peptide genes (*ang* 1, 3, and 4). However, only *ang4* is highly expressed in the colon and small intestine and is produced by goblet and Paneth cells [68–70]. Humans only have one orthologous protein (ANG). Both *ang4* and ANG exhibit antifungal activity *in vitro* [68]. Similarly, CX3CR1 mainly expressed by macrophages in the intestine plays a critical role in antifungal defense to CA [55, 71]. Furthermore, recent evidence indicates that CX3CR1 expressing macrophages are important to maintain intestinal barrier function to prevent bacterial translocation from the gut [72]. Mice dosed with 1% TCA orally had significantly decreased abundance of *L. johnsonii, T. sanguinis*, and *C. celatum* in the cecal contents and colon mucosal scrapings. Interestingly, previous findings from our lab indicates that all these three bacterial genera were also decreased in antibiotic-treated mice susceptible to CA [25]. Commensal bacteria such as *L. johnsonii* plays a critical role in maintaining the intestinal barrier function [73–76] and can directly inhibit the growth of fungi [77]. Since *ang4* and *Cx3cr1* expression is also regulated by the microbiota [68, 78], it is possible that TCA indirectly regulates CA by controlling *ang4* and *Cx3cr1* expression through one of the three commensal bacteria modulated by TCA.

In the intestine, TCA is mainly absorbed through apical sodium-dependent bile acid transporter (ASBT), encoded by the *Slc10a2* gene in ileal and colonic epithelial cells [49, 57, 58]. After absorption, TCA binds to FXR, a nuclear receptor encoded by *Nr1h4*, expressed in enterocytes and macrophages resulting in the transcription of target genes [49, 57, 59–61]. The expression of *Slc10a2* and *Nr1h4* was also significantly increased in the colon of antibiotic treated mice [49, 79]. Therefore, these findings along with others [55, 68, 71, 72] suggest that TCA may induces CA colonization and dissemination by controlling intestinal defense system and barrier function through ASBT and FXR. Future studies to understand how TCA impacts these factors in regulating CA will form a strong platform to reduce fungal colonization and dissemination from the GI tract.

Under normal physiological conditions, intestine and cecum holds a bile salt concentration gradient ranging from 2% and 0.05% [80, 81], therefore we used an average concentration of 1% TCA for mouse studies. Therefore, future studies to understand the effect of TCA at different concentrations and dose will gain insights into the role of TCA on CA colonization and dissemination. For example, a lower concentration (0.1 or 0.5%) may affect only microbiota or host and potentially can enhance only CA colonization without dissemination. Further, in order to rule the possibility that the metabolic transformation of orally administered TCA to DCA may contribute to host and microbiota alterations observed in our study, we also examined the DCA levels in TCA treated mice. Surprisingly, mice that received TCA had considerably reduced levels of DCA compared to untreated groups (Supplementary Figure 5). This may be because TCA may directly inhibit certain species of bile-metabolizing bacteria (*Lactobacillus*). Previous findings from our lab indicates that TCA promotes, whereas DCA inhibits, CA growth *in vitro* [25, 82, 83]. However, surprisingly our *in vivo* findings indicate that orally administration with TCA alone in mice induces fungal colonization and dissemination suggesting that regulating host and microbiota is important in order to cause intestinal barrier dysfunction leading to CA dissemination and mortality [5, 53, 74, 84]. In addition to host and microbiota effect, TCA and (or) DCA may have a direct effect on CA. Future studies warrants further investigation on the direct effect of TCA and DCA on CA, and the *in vivo* role of DCA on fungal colonization and dissemination.

Given the significant alteration in bile acid levels during various therapies and diseases [29–51, 85–106], understanding how bile acids regulate CA will provide a new approach for to manipulate and control the fungal-bile communication system in favor of the host. Furthermore, CA in the gut plays an active role in inflammatory bowel disease [95, 116, 118], the microbiota-gut-brain axis [110, 119], and treating various pathogens including *Clostridium difficile* [120, 121]. Our findings will have a significant impact in understanding how microbial metabolites interact with the microbiota and host in regulating gut fungi.

Given the high morbidity and mortality associated with invasive fungal infections, the limited arsenal of antifungals, and the continuing emergence of treatment-resistant strains, new approaches to treating and preventing invasive fungal infections in patients are desperately needed. Though TCA and DCA are well-known metabolites, their specific role in the colonization and pathogenesis of CA is unknown. Addressing this critical gap in our knowledge could pave the way to use these naturally occurring bile acid metabolites to reduce fungal colonization and ultimately decrease dissemination in the host. In depth understanding of the role of bile acid metabolites on fungal colonization and dissemination will form a strong platform to modulate bile acids or bile-acid metabolizing commensal bacteria or to modulate the host defense system through bile acids to prevent and treat invasive fungal infections originating from the GI tract.

