## Supplementary Materials for "Bile Acid Regulates the Colonization and Dissemination of *Candida albicans* from the Gastrointestinal Tract by Controlling Host Defense System and Microbiota"

**Supplementary Figure 1.** Bar graph showing the alpha diversity from cecal contents and mucosal scrapings from colon in the untreated and TCA treated mice groups infected with CA.

**Supplementary Figure 2-4.** Heatmap showing the relative abundance of bacterial members (phylum, order and family) from cecal contents and mucosal scrapings from colon in the untreated and TCA treated mice groups infected with CA.

**Supplementary Figure 5.** TCA and DCA levels in the cecal contents from untreated and TCA treated mice infected with CA collected after 10-12 days of infection and treatment were shown here. 4-5 mice per group was used and the data represents mean ± SEM. Statistical significance was evaluated using the Student's t test. *p* values ≤0.05 (*) or ≤0.01 (**) were considered statistically significant.

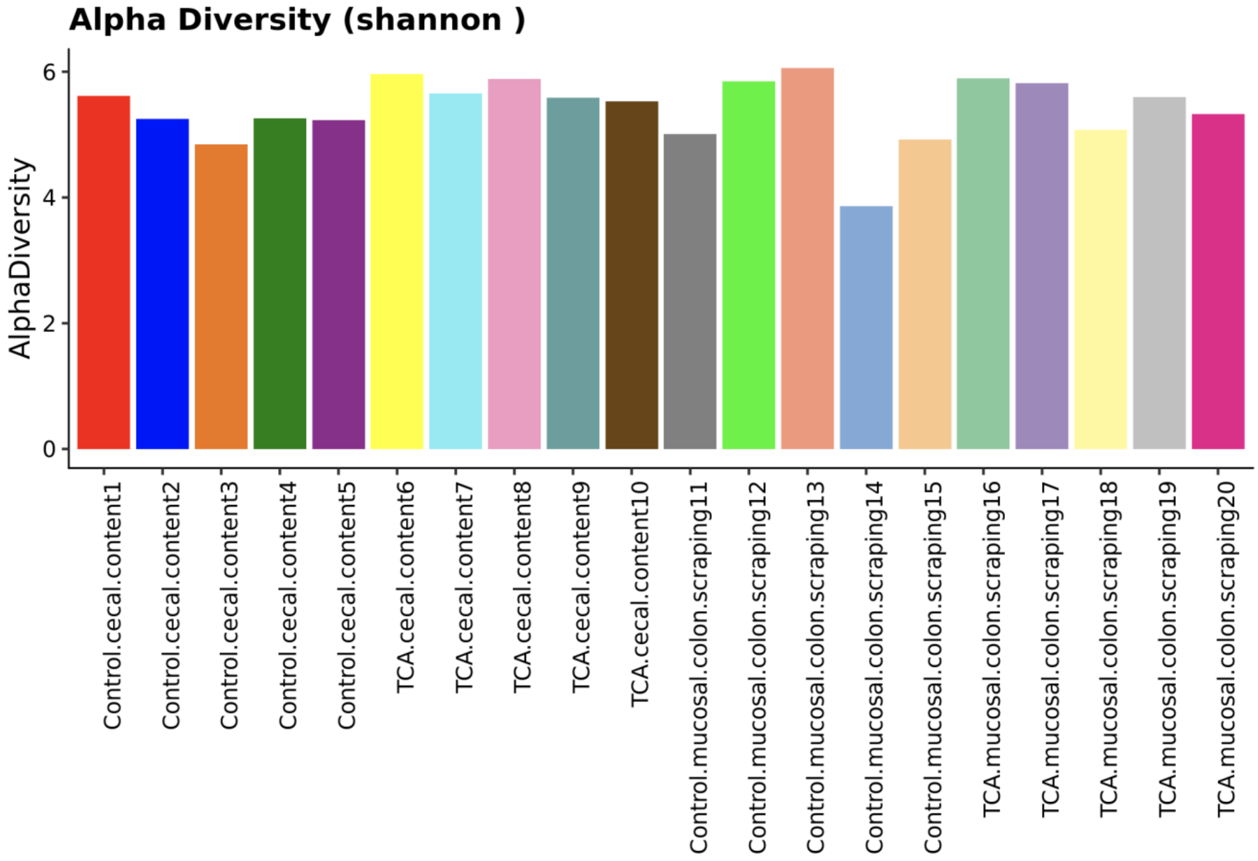


**Supplementary Figure 1.**

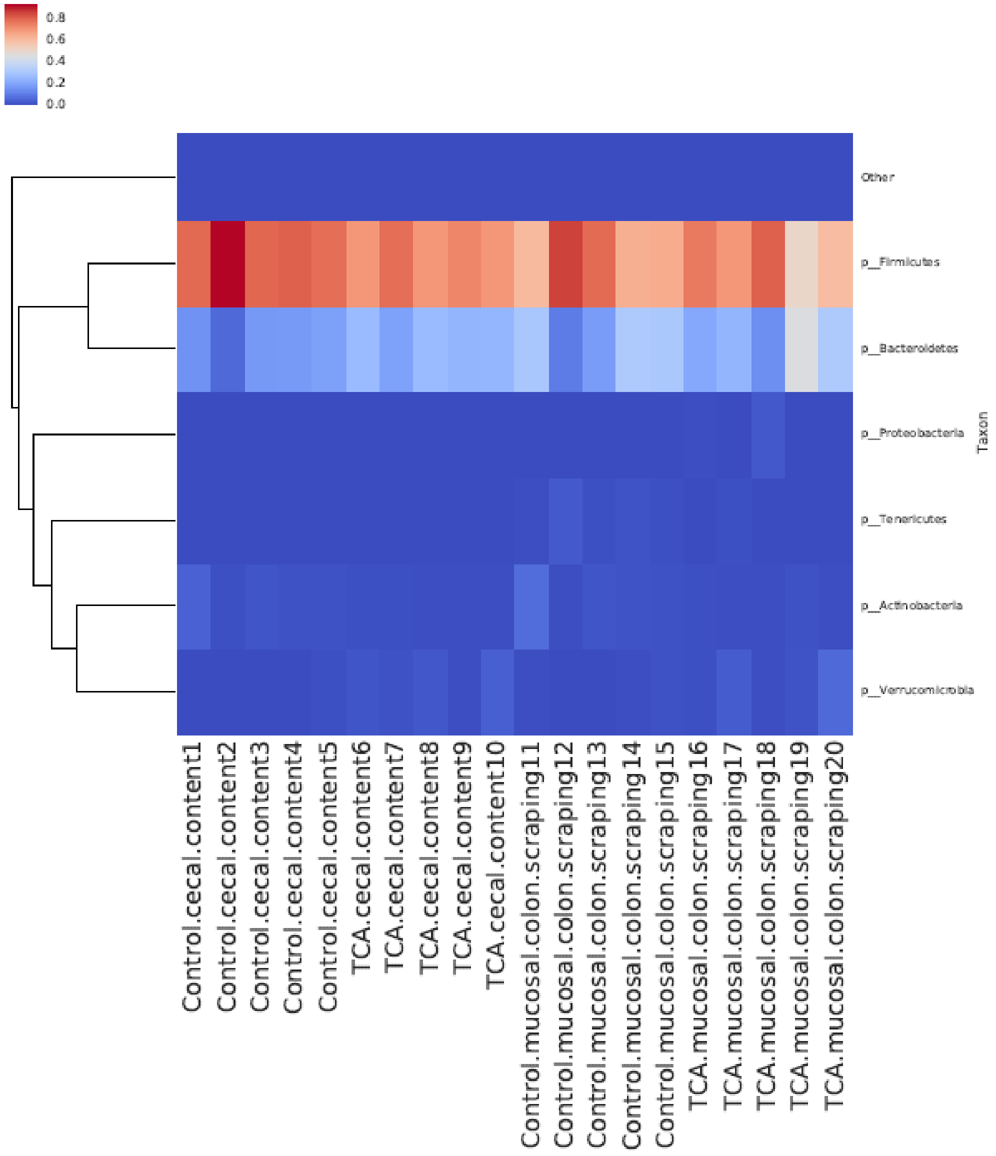


**Supplementary Figure 2.**

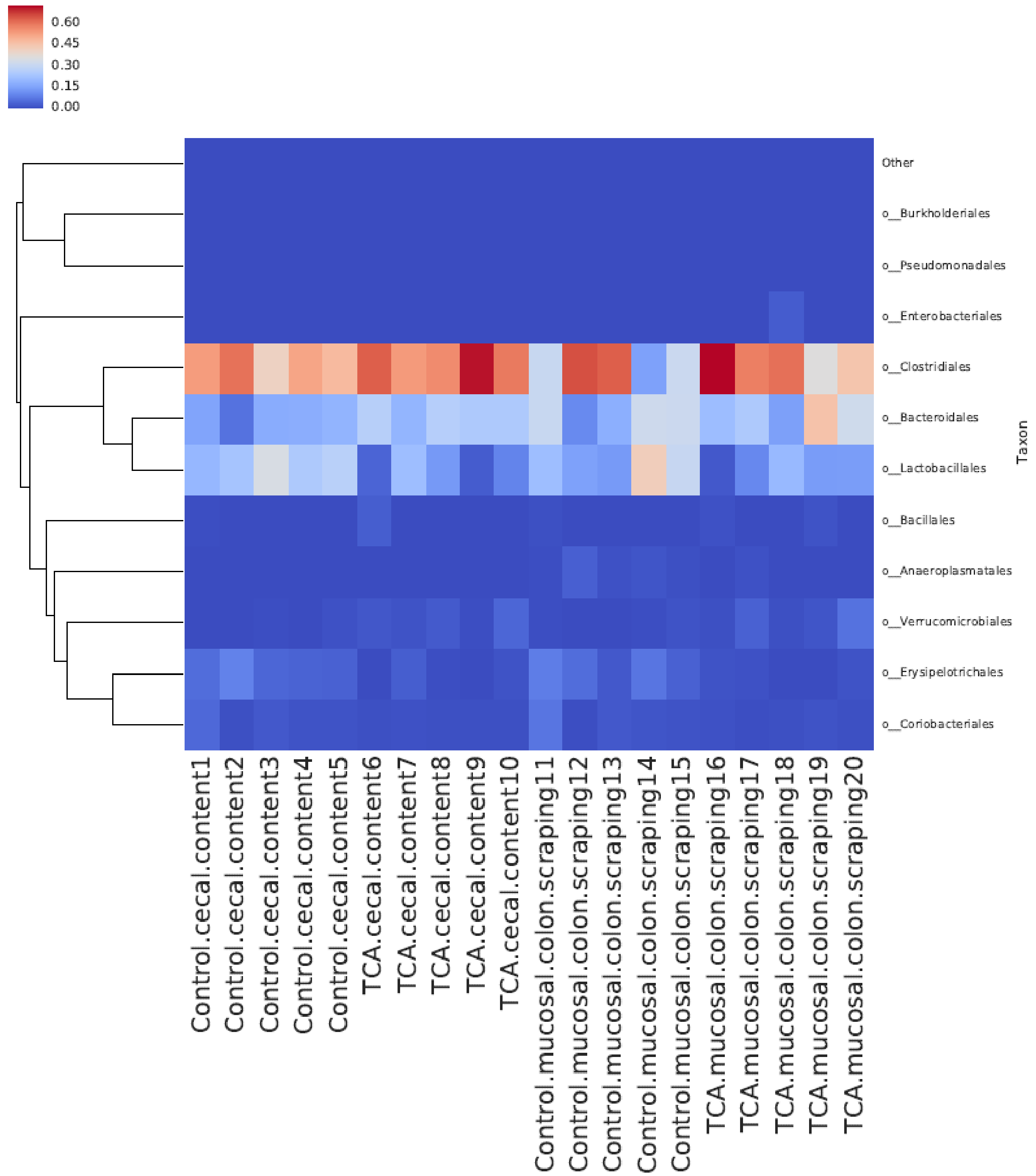


**Supplementary Figure 3.**

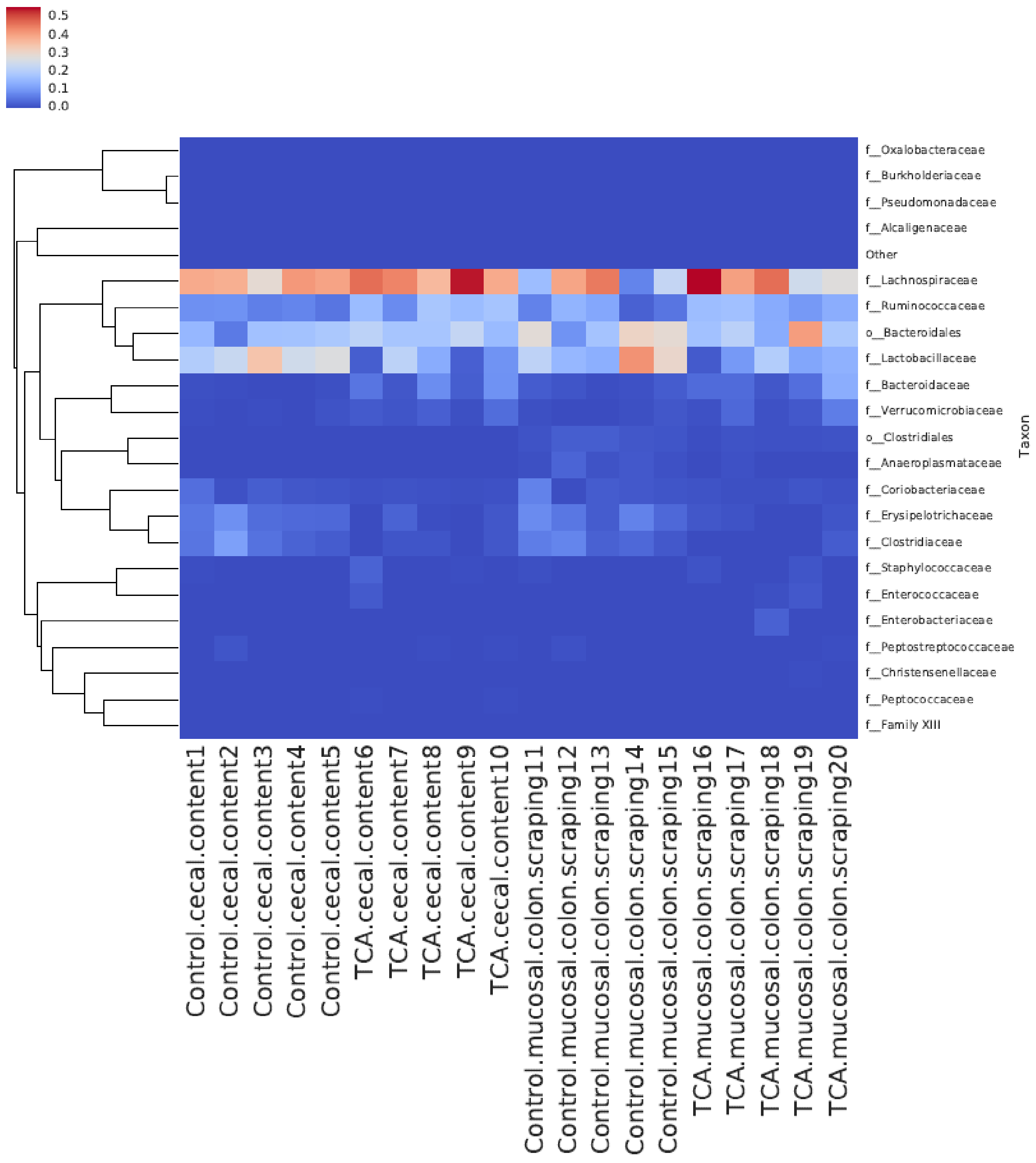


**Supplementary Figure 4.**

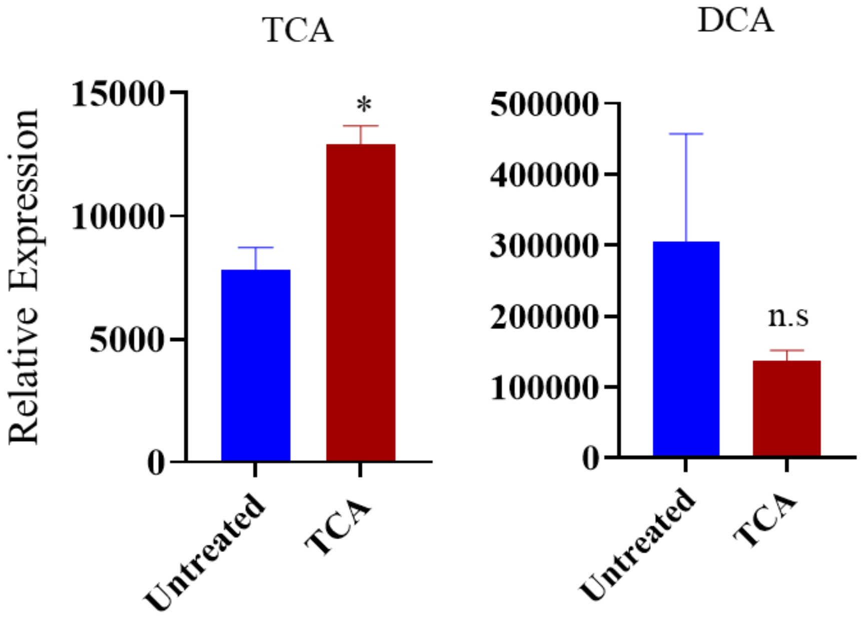


**Supplementary Figure 5.**
